# AAV serotype PHP.eB achieves superior neuronal transduction efficiency compared to AAV9 in pigtail macaques following intracerebroventricular administration

**DOI:** 10.1101/2025.02.04.636515

**Authors:** Michal G. Fortuna, Lyle Nyberg, Naz Taskin, Natalie Weed, Avery Hunker, Kathryn Gudsnuk, Melissa Berg, Boaz Levi, Ed Lein, Jonathan T. Ting

## Abstract

Adeno-associated virus (AAV) vectors are pivotal in gene therapy for neurological disorders due to their ability to enable long-term gene expression in the central nervous system (CNS). However, transducing larger brains, such as those of non-human primates (NHPs), remains challenging, necessitating alternative delivery routes and optimized capsids. This study directly compares the transduction efficiency and biodistribution of the benchmark AAV9 and its engineered derivative, AAV-PHP.eB, following intracerebroventricular (ICV) administration in juvenile *Macaca nemestrina*. Employing a neuron-specific promoter and nuclear-localized reporter, we systematically quantified transduction across cortical, subcortical, and spinal regions. AAV-PHP.eB demonstrated significantly higher transduction rates in cortical and spinal regions compared to AAV9, despite similar expression patterns. Both vectors exhibited limited subcortical penetration and significant peripheral leakage, highlighting key challenges in CNS targeting. This is the first study to quantitatively compare AAV-PHP.eB and AAV9 in NHPs, providing valuable insights into the advantages and limitations of engineered AAV capsids for CNS gene therapy. These findings lay a critical foundation for optimizing vector designs and delivery strategies to improve outcomes in clinical applications for neurodegenerative and neurodevelopmental disorders.

## I. Introduction

Neurological diseases, such as Parkinson’s, Alzheimer’s, and various inherited disorders have long posed significant challenges for effective treatment, primarily due to the complex nature of brain physiology, and the blood-brain barrier (BBB), which limits access of systemically delivered therapeutic agents to the central nervous system (CNS). Gene therapy offers an alternative in treating neurological disorders by correcting underlying genetic defects or delivering therapeutic transgene cargos to restore normal brain function (1,2). Adeno-associated viruses (AAVs) have become pivotal tools for gene therapy due to their ability to deliver genetic material safely and with sustained expression (1,3,4).

Among all available AAV serotypes, AAV9 has been widely adopted in both preclinical and clinical studies to address CNS disorders due to its ability to traverse the BBB and transduce a broad range of CNS cell types, including neurons and glia (3–5). However, AAV9 has relatively low BBB crossing efficiency in large primate brains, necessitating high viral loads when delivered intravenously (IV), raising safety concerns due to potential immune responses and off-target effects (6,7).

Extensive efforts have been made to engineer AAV capsids to improve their tropism and biodistribution profile and explore alternative delivery routes and methods (1,3,4,8). One promising approach is intracerebroventricular (ICV) administration, which bypasses the need for BBB crossing by delivering AAVs directly into the cerebrospinal fluid (CSF). This method, in principle, allows for a larger distribution of viral particles across the CNS, facilitating broader transduction compared to intravenous or intrathecal delivery (6,9,10)

AAV-PHP.eB is an engineered variant of AAV9 capsid with enhanced CNS tropism and superior transduction efficiency following IV delivery (8,11). However, these effects are species-specific as non-human primates (NHPs) lack LY6A (SCA-1), the receptor for BBB crossing in rodents (12). Nevertheless, a recent study in adult rhesus macaques showed that AAV-PHP.eB achieved greater CNS transduction than its predecessor, AAV-PHP.B, when delivered via intrathecal (IT) administration (13). The efficacy of AAV-PHP.eB following ICV administration in NHPs remains untested, leaving open the question of whether it offers superior transduction compared to AAV9 in this delivery context. Using ICV delivery of clinically relevant doses in juvenile *Macaca nemestrina* and a neuron-specific, nuclear-localized reporter, we systematically quantified neuronal transduction across the brain and spinal cord and show that AAV-PHP.eB achieves superior CNS transduction vs. AAV9.

## II. Materials and Methods

### A. Experimental design and surgery

#### Non-human primates

Experiments with macaque monkeys conformed to the guidelines provided by the US National Institutes of Health and were approved by the University of Washington Animal Care and Use Committee. The experiments reported herein were performed on seven juvenile male *Macaca nemestrina* monkeys weighing 3.05-4.5kg. All animals used in this study were pre-screened and determined to be seronegative for AAV9 neutralizing antibodies. Three animals received AAV-PHP.eB virus injection (4.4-4.5kg), three received AAV9 injection (3.05-3.35kg) and one control subject received saline injection (4.5kg). Virus injected animals received a dose of 4e13 vg/kg of respective virus (1.2-1.8ml total infusion volume).

#### Viral vectors

Both serotypes carried the same transgene cassette, namely 3xFLAG tagged Histone 2B under the control of the human Synapsin 1 promoter. AAV9 and AAV-PHP.eB viral particles were produced in HEK293 cells by transient transfection with plasmids containing the transgene cassette, helper plasmid, and rep/cap sequences (14). The clarified cell lysate was purified using POROS CaptureSelect AAVX affinity chromatography, followed by ultracentrifugation in a cesium chloride density gradient to enrich for full capsids (15,16). The collected AAV particles were concentrated and buffer-exchanged using a centrifugal filter unit with a 100 kDa molecular weight cutoff. The final AAV preparation was aliquoted and stored at −80°C until further use. The viral titer was determined by measuring the genome copies using qPCR with transgene-specific primers.

#### Surgery and intracerebroventricular administration

Viral vectors were injected during surgery in an operating room under isoflurane and fentanyl anesthesia. Planning of the injection and corresponding stereotactic coordinates were based on prior anatomical MRI scans of each subject. Once anesthetized, subjects were positioned in the stereotactic frame, skull was exposed, and a burr hole was drilled few millimeters off the midline at the predetermined AP and ML levels. A needle was mounted on the stereotactic arm, positioned over the opening, and dura was nicked at the site of tip insertion. The needle was lowered through the opening to planned depth and correct placement in the ventricle was verified by drawing a small amount of CSF into the syringe. Once access to the ventricle was confirmed, virus solution was loaded into a syringe. Solution was manually infused as follows: total virus load (1.2-1.8ml) was divided into six equal parts (0.2-0.3ml per part), and each part was injected over 10 min. Thus, injection of the whole volume lasted about 1h for each animal. After completing the injection, the needle was removed, and the brain surface rinsed with sterile saline. A piece of DuraGen (Integra, USA) was fitted into the burr hole to cover the opening. The subcutaneous tissues and skin were sutured closed, tissue adhesive (3M Vetbond Tissue Adhesive 1469) was applied, and incisional bupivacaine was administered. After the surgery, animals were placed in individual cages overnight and resocialized after recovery. Surgery and animal husbandry was provided by the staff of the Washington National Primate Research Center (WaNPRC).

### B. Tissue processing

Six weeks post-surgery, animals were euthanized with pentobarbital and transcardially perfused with NMDG artificial CSF (NMDG aCSF) solution (17). The brain and spinal cord (sectioned into cervical, thoracic, and lumbar/sacral segments) were collected; spinal segments were fixed in 10% neutral-buffered formalin (NBF). The brain was placed in ice-cold NMDG aCSF for transport, then hemisected, and sliced into ∼5mm coronal slabs (12–14 per hemisphere). Alternate slabs were NBF-fixed (48–72h) or flash-frozen in isopentane (17). Fixed slabs underwent 30% sucrose infiltration and sectioning into 50 µm coronal slices on a sliding microtome (Leica SM2000R). Sections were stored in PBS with sodium azide at 4 °C for IHC processing. Peripheral organs samples (liver, kidney, muscle, heart) were snap-frozen in liquid nitrogen, and stored at −80°C until further use.

### C. Immunohistochemistry

Complete immunostaining procedure on free-floating sections was described previously (18). The following primary antibodies were used: (1) anti-FLAG (1:500, mouse, M2 from Sigma), (2) anti-NeuN (1:1000, rabbit, ABN78 from Millipore), (3) anti-Parvalbumin (1:5000, mouse, PV235 from Swant), (4) anti-Calretinin (1:2000, rabbit, Cat# CR7697 Swant), (5) anti-Iba1 (1:1000, rabbit, Cat# 019-19741, Wako), (6) anti-Glut1 (1:1000, rabbit, Cat# 07-1401 Millipore). Secondary antibodies included (all from Invitrogen): (1) Alexa 555 donkey anti-rabbit (1:500, A32749), (2) Alexa 568 donkey anti-rabbit (1:500, A10042 or A10042), (3) Alexa 488 goat anti-mouse (1:500, A11029), (4) Alexa 647 donkey anti-mouse (1:1000, AB_162542). DAPI (Cat# 62248 Thermo Scientific™) was added at 1-2ug/mL for nuclear counterstain.

Spinal Cord segments were paraffin embedded post fixation, cut into 5µm sections and mounted onto glass slides. For IHC, standard deparaffinization and rehydration protocol was used (Xylene and Ethanol gradient), followed by antigen retrieval using heat induced step: 10min wash with TBS at RT, 30min incubation in CC1 reagent (Roche, product code 950-124) at 95°C, and washed 3 x 5min in TBS at RT. After antigen retrieval, standard IHC protocol for FLAG and NeuN staining was followed, with modified Ab concentrations: 1:3000 for FLAG and NeuN primary Ab, and 1:2000 for secondary Ab.

### D. Molecular biology

For DNA isolation, frozen tissue was moved from −80° to −20° an hour before tissue punching (with ‘Disposable Biopsy Punch’, Robbins Instruments, Ref# RBP-40), which was done on a dissection table chilled to −20°. Viral DNA was isolated with DNeasy® Blood & Tissue Kit (Qiagen). After isolation, DNA concentration was measured (NanoDrop, Thermo Scientific™), and diluted to working concentrations ranging from 0.03-10ng/ul. Prior to PCR reaction, template DNA was treated with SmaI restriction enzyme (New England Biolabs). PCR was performed with PTC Tempo Deepwell Thermal Cycler (Bio-Rad). AAV genome copies were quantified by ddPCR in ddPCR Supermix for Probes (Bio-Rad —no. 18630254) with WPRE3 primer/probe (Bio-Rad – no. 10042959) for viral DNA, and MRFAP1 was used as a reference gene (Fw 5’-GAACCGGAGCAAGCTGT-3’, Rv 5’-GGTTCTGGAGGTGGTTGAG-3’, probe 5’-/5HEX/-CACCTGCGTTTTGATCTGGATGAGC/3IABkFQ/-3’; from Integrated DNA Technologies, Inc.). Droplets were generated with the QX200 Droplet Generator following the manufacturer’s instructions (Bio-Rad). Droplets were read with the QX200 Droplet Reader and results were analyzed with QuantaSoft Analysis Pro Software (V1.0.596-Bio-Rad).

### E. Microscopy, image processing and cell counting

Fluorescent images were taken with either an Axio Observer 7 epifluorescent microscope controlled by ZEN 2.3 (blue edition, Zeiss, Jena, Germany) or Nikon Ti-Eclipse 2 epifluorescent microscope and further processed with ImageJ. Standard image adjustments, including pseudo-coloring, thresholding, brightness, and contrast adjustments, were applied equally across the entire image.

For quantitative analysis, five cortical areas were selected: prefrontal (vmPFC), primary motor (M1), somatosensory (S1/S2), parietal (PEC and PEa (MIP)), and visual (V1/V2) cortices. For each investigated region, at least three different sections were imaged, and at least three different ROIs were selected and quantified per image (for statistical comparison, multiple ROIs from the same section were averaged resulting in one value per slice). Standard ROI was a full cortical column 1-1.5mm wide and 2-3mm deep (average of 4,046 NeuN cells per ROI).

Images were processed with ImageJ. NeuN and FLAG channels were handled separately. Cells were quantified using the Analyze Particles tool. The neuronal transduction rate was calculated as a fraction of FLAG^+^ cells to NeuN^+^ cell number.

### F. Statistics

For statistical comparison SigmaPlot (Grafiti LLC, Palo Alto, CA) was used. Results are expressed as a mean (or median) ± standard error. To compare 2 groups t-test was used (normality test: Shapiro-Wilk; Mann-Whitney Rank Sum Test if data was not normally distributed). One Way ANOVA was used to compare multiple groups (Kruskal-Wallis One Way Analysis of Variance on Ranks) followed by All Pairwise Multiple Comparison Procedures (Dunn’s Method). Differences were considered significant if p < 0.05.

## III. Results

To facilitate comparison of neuronal transduction and biodistribution of AAV9 and AAV-PHP.eB in the NHP brain, we designed a viral vector to label neuronal nuclei ubiquitously across the brain. The vector was packaged into both AAV9 and PHP.eB capsids and unilaterally injected ICV at a dose of 4 × 10¹³ vg/kg in juvenile *Macaca nemestrina* (**Figure 1A**). The administered viral load was well tolerated in all subjects, with no noticeable physiological or behavioral abnormalities observed post-injection. After six weeks, the brain, spinal cord, and peripheral organs were collected and processed for molecular and histological analysis (**Figure 1B**).

**Figure 1.**
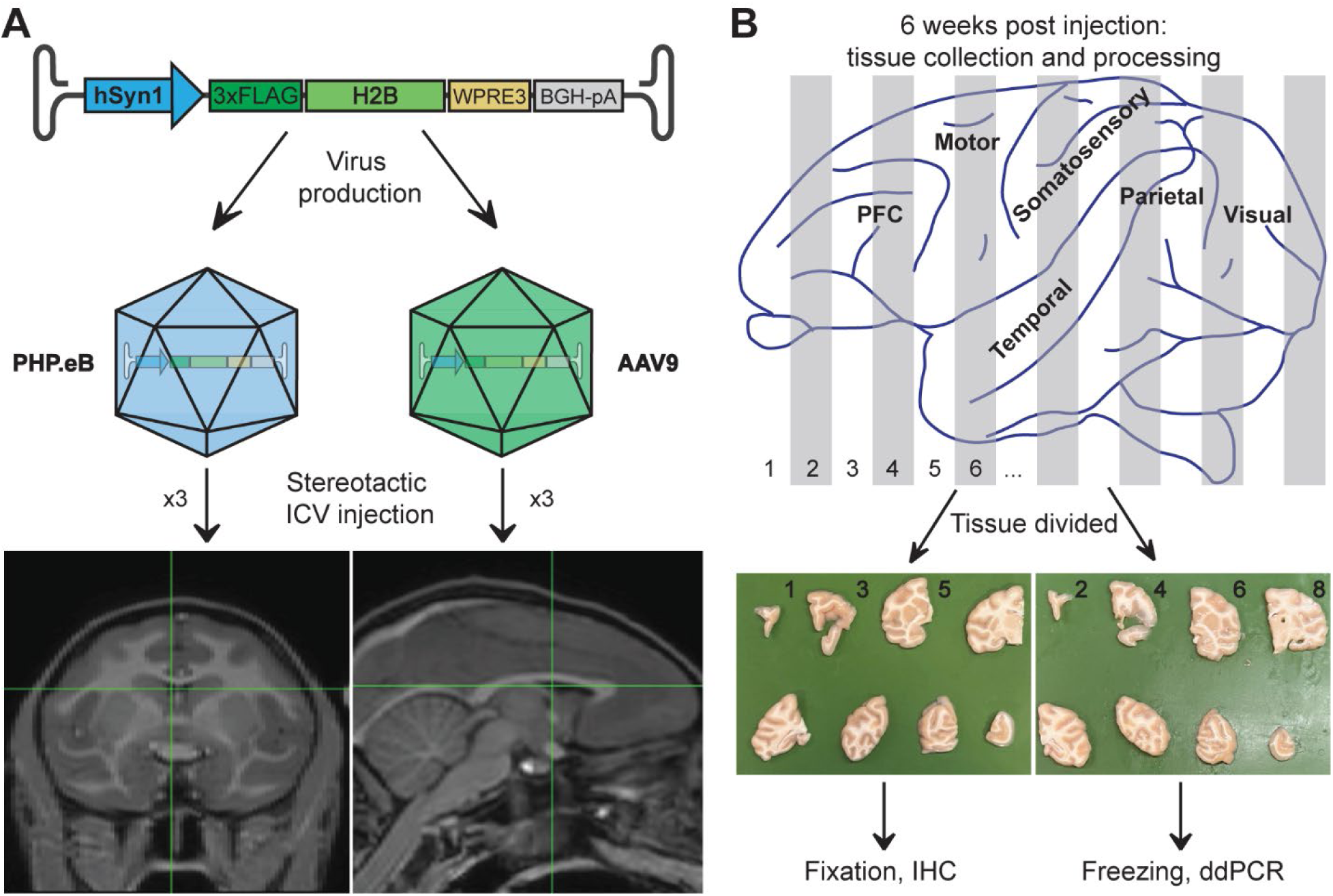
Experimental design and tissue processing. **A)** Schematic of the DNA construct used for packaging of AAV-PHP.eB and AAV9 vectors, which were unilaterally injected into the lateral ventricle of 3 animals per group. MRI images show example coordinates of the target injection site in one of the experimental subjects. The DNA construct includes: hSyn1 – the human synapsin 1 gene promoter; 3xFLAG tag fused with Histone 2B (H2B) for nuclear localization; WPRE3 – Woodchuck Hepatitis Virus Posttranscriptional Regulatory Element; BGH-pA – bovine growth hormone polyadenylation signal. Hairpin loops represent inverted terminal repeats (ITRs) flanking the expression cassette. **B)** Schematic of downstream tissue processing: brains were cut into ∼5 mm coronal slabs, and alternate slabs were either formalin fixed for immunohistochemistry (IHC) or frozen for genetic material extraction for ddPCR analysis. Major areas of the cerebral cortex are indicated in the drawing; PFC-the prefrontal cortex.

### Unilateral ICV delivery results in widespread CNS transduction

We first qualitatively evaluated the relative expression profiles for AAV-PHP.eB vs. AAV9 vectors by immunostaining for the 3xFLAG tag under matched staining and imaging conditions. Despite performing unilateral injections, we observed no obvious difference in number of FLAG^+^ cells between the ipsilateral and contralateral hemispheres across both groups (**Figure 2**). Both vectors labeled the brain parenchyma to varying degrees of completeness, ranging from occasional nuclei and sparse cell groupings to densely transduced areas resulting in irregular coverage. Most FLAG^+^ nuclei were observed in the cerebral cortex. Areas in the vicinity of the injection site or in direct contact with the lateral ventricle were typically more densely labeled. In contrast, deeper, subcortical structures such as putamen and thalamus regions more distant from the site of CSF delivery showed only a few FLAG^+^ cells scattered throughout (**Figure 2** and **3B**). Although both serotypes yielded a similar overall pattern of transgene expression, there was an obvious difference in the number of FLAG^+^ cells, with the PHP.eB cohort exhibiting higher labelling density compared to AAV9. Despite individual variations in expression pattern across the brain (see **Table 1**), there was a general trend toward more intense labeling in the posterior cortices (e.g., parietal, and visual cortex) and less prominent viral penetration in the temporal lobe and prefrontal areas (**Figures 2 A2/3 and B2/3**). The motor and somatosensory cortices were consistently among the areas with the highest levels of transduction.

**Figure 2.**
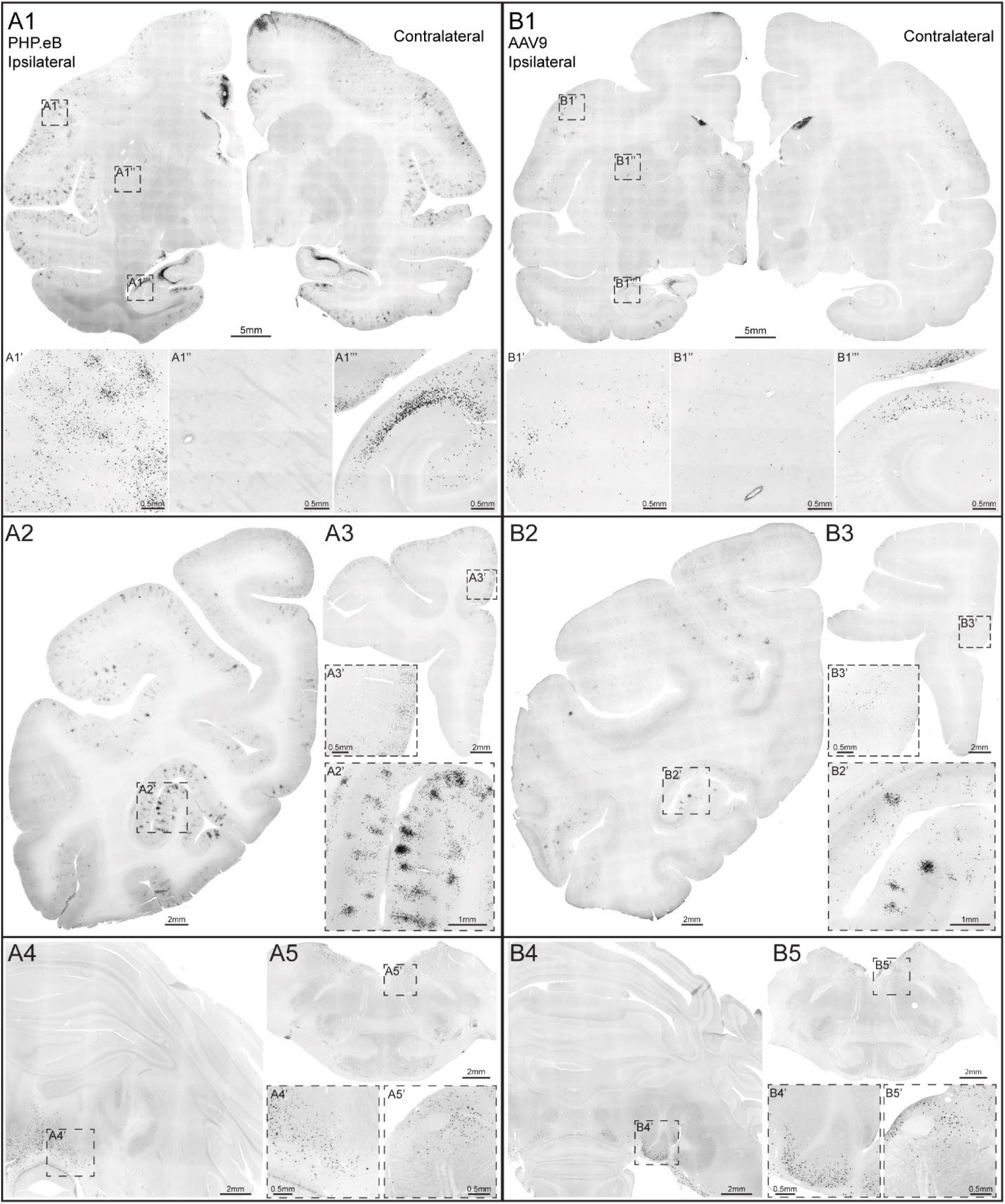
Unilateral injection of PHP.eB and AAV9 results in brain-wide transgene (FLAG) expression. **A1/B1**: Representative coronal slices of the left and right hemispheres from a level near the injection site. **A1**: PHP.eB, **B1**: AAV9. Dashed boxes indicate the locations of zoomed-in images shown at the bottom of the figure: **A1’/B1’** – ventral premotor cortex, **A1’’/B1’’** – putamen, **A1’’’/B1’’’** – hippocampus. **A2/3 and B2/3**: Representative coronal sections at the level of the occipitoparietal (**A2/B2**) and prefrontal (**A3/B3**) lobes; dashed boxes show the locations of zoomed-in areas. **A4/B4**: Example of cerebellum; note the lack of expression in the cerebellar cortex, with expression only in the deep cerebellar nuclei (**A4’/B4’**). **A5/B5**: Example coronal section of the brainstem; zoomed-in images of the area adjacent to the abducens nucleus (6N) and the 4^th^ ventricle (**A5’/B5’)**. Scale bars are shown in each micrograph.

**Figure 3.**
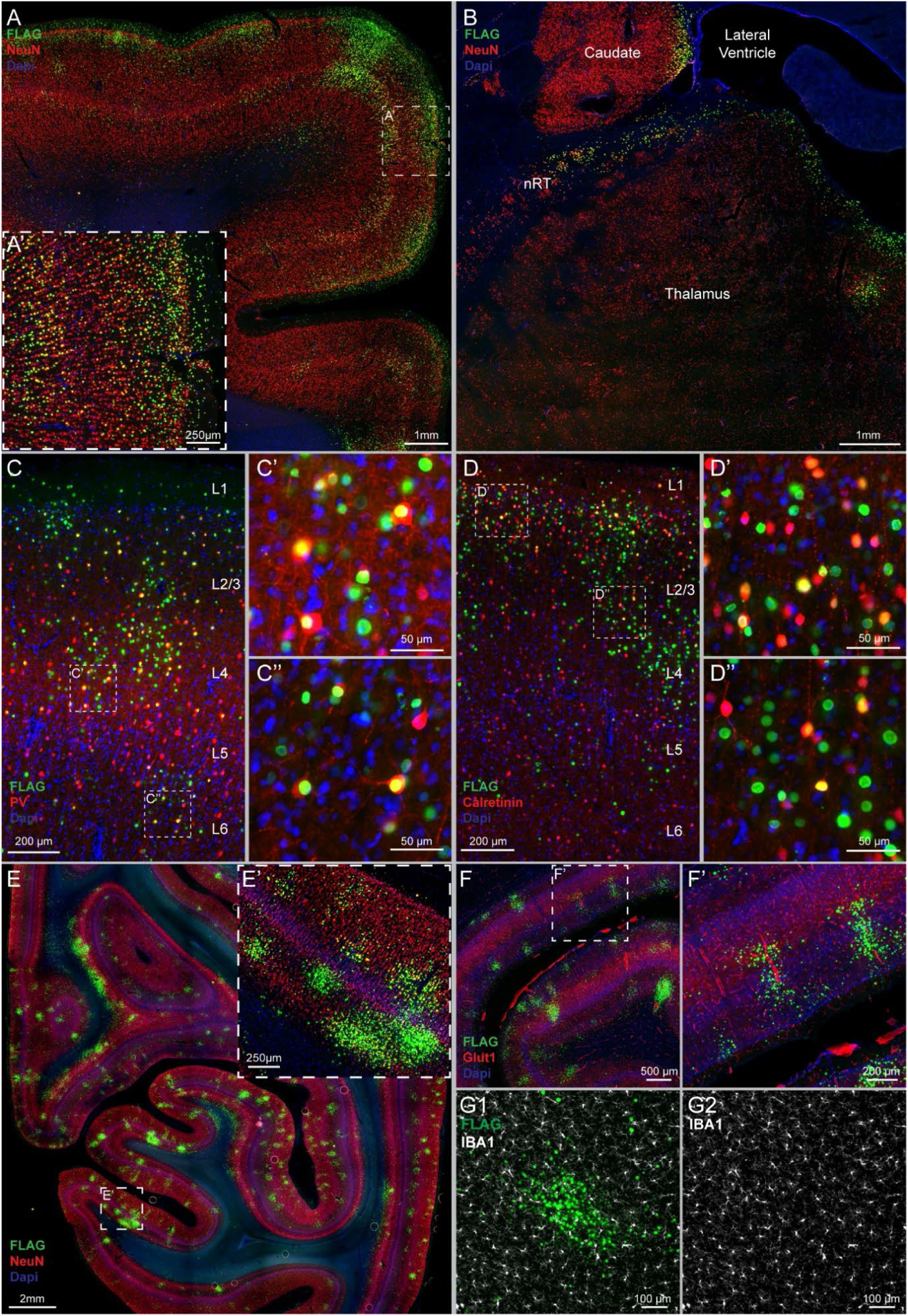
Expression pattern across the brain after ICV vector delivery. **A and B)** Examples of expression patterns in cortical and subcortical structures near the injection site: **A** – supplementary motor area; **B** – anterior caudate and thalamus bordering the lateral ventricle (nRT – thalamic reticular nucleus). **C**: Co-expression of transduced cells with Parvalbumin, a marker for a subpopulation of inhibitory interneurons. **D**: Co-expression of transduced cells with Calretinin, another marker for a subpopulation of GABAergic interneurons. In **C** and **D**, dashed boxes mark the locations of zoomed-in sites, shown in **C’/C’’** and **D’/D’’**; approximate cortical layer locations are indicated. **E**: Example of uneven, patchy labeling in the occipital cortex; **E’** – magnification of the area boxed in **E**. **F**: Colocalization of some transgene expressing cell clusters with larger blood vessels, revealed by Glut1 co-staining; **F’-** magnification of the area boxed in **F**. **G1**: Overlay of a cluster of FLAG^+^ nuclei with microglia cells, stained with IBA1; **G2 –** IBA1 signal. A pseudo-coloring scheme and scale bars are shown in each micrograph.

**Table 1.**
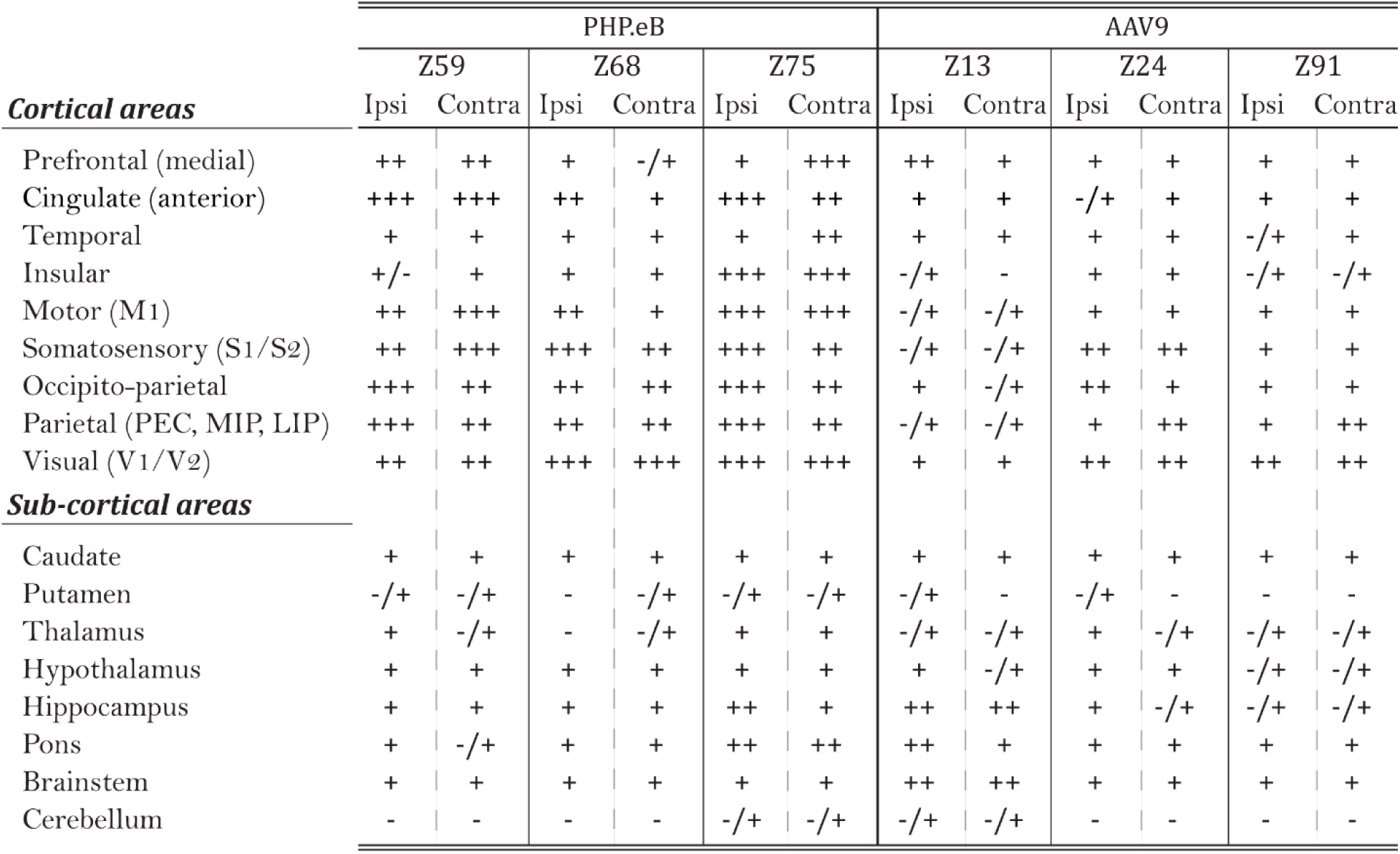
Qualitative analysis of transgene expression throughout the brain in each subject. . Selected cortical and subcortical areas were examined for the degree of transduction. The rate of transduction was scored as follows: **-/+**: sparse labeling, individual cells; **+**: low expression (<5% NeuN); **++**: medium expression (distributed cells and occasional cell clusters, ∼5-10% NeuN); **+++**: high expression (densely labeled neuropil with numerous cells and frequent cell clusters, >15% NeuN). **Abbreviations**: M1 - primary motor cortex, S1/S2 - somatosensory area 1 and 2, PEC - parietal area PE caudal part, MIP – medial intraparietal area, LIP - lateral intraparietal area, V1-primary visual cortex, V2 – visual area 2.

|  | PHP.eB |  |  |  |  |  | AAV9 |  |  |  |  |  |
| --- | --- | --- | --- | --- | --- | --- | --- | --- | --- | --- | --- | --- |
|  | Z59 |  | Z68 |  | Z75 |  | Z13 |  | Z24 |  | Z91 |  |
|  | Ipsi | Contra | Ipsi | Contra | Ipsi | Contra | Ipsi | Contra | Ipsi | Contra | Ipsi | Contra |
| **Cortical areas** |  |  |  |  |  |  |  |  |  |  |  |  |
| Prefrontal (medial) | ++ | ++ | + | -/+ | + | +++ | ++ | + | + | + | + | + |
| Cingulate (anterior) | +++ | +++ | ++ | + | +++ | ++ | + | + | -/+ | + | + | + |
| Temporal | + | + | + | + | + | ++ | + | + | + | + | -/+ | + |
| Insular | +/- | + | + | + | +++ | +++ | -/+ | - | + | + | -/+ | -/+ |
| Motor (M1) | ++ | +++ | ++ | + | +++ | +++ | -/+ | -/+ | + | + | + | + |
| Somatosensory (S1/S2) | ++ | +++ | +++ | ++ | +++ | ++ | -/+ | -/+ | ++ | ++ | + | + |
| Occipito-parietal | +++ | ++ | ++ | ++ | +++ | ++ | + | -/+ | ++ | + | + | + |
| Parietal (PEC, MIP, LIP) | +++ | ++ | ++ | ++ | +++ | ++ | -/+ | -/+ | + | ++ | + | ++ |
| Visual (V1/V2) | ++ | ++ | +++ | +++ | +++ | +++ | + | + | ++ | ++ | ++ | ++ |
| **Sub-cortical areas** |  |  |  |  |  |  |  |  |  |  |  |  |
| Caudate | + | + | + | + | + | + | + | + | + | + | + | + |
| Putamen | -/+ | -/+ | - | -/+ | -/+ | -/+ | -/+ | - | -/+ | - | - | - |
| Thalamus | + | -/+ | - | -/+ | + | + | -/+ | -/+ | + | -/+ | -/+ | -/+ |
| Hypothalamus | + | + | + | + | + | + | + | -/+ | + | + | -/+ | -/+ |
| Hippocampus | + | + | + | + | ++ | + | ++ | ++ | + | -/+ | -/+ | -/+ |
| Pons | + | -/+ | + | + | ++ | ++ | ++ | + | + | + | + | + |
| Brainstem | + | + | + | + | + | + | ++ | ++ | + | + | + | + |
| Cerebellum | - | - | - | - | -/+ | -/+ | -/+ | -/+ | - | - | - | - |

Mostly sparse or area-specific labeling could be observed throughout the hypothalamus (e.g., paraventricular hypothalamus), parts of the midbrain (e.g., the periaqueductal gray, superior colliculus), and various nuclei of the hindbrain (e.g., pontine nuclei, hypoglossal nucleus). Hindbrain structures closer to the fourth ventricle and medullary surface showed considerable transduction, whereas very few FLAG^+^ cells could be found in core of the tissue. The cerebellum, aside from aspects of the deep cerebellar nuclei (DCN) neighboring the fourth ventricle, showed no appreciable transgene expression (**Figure 2 A4/5 and B4/5**). Table 1 summarizes qualitative analysis of major brain areas for all subjects.

Detailed comparative examination showed that both serotypes displayed similar characteristics in terms of resultant expression pattern, where both groups lacked uniform transduction. For instance, in the context of the same anatomical structure, a region with high density of FLAG^+^ cells could neighbor areas displaying very few FLAG^+^ cells (**Figure 3A**). Additionally, there appeared to be a tendency for labeled cells to occupy the upper cortical layers 2/3 or deeper layer 5 (**Figure 3A/A’, 3E**), suggesting that excitatory pyramidal neurons may be preferentially transduced. To explore this further, we co-stained the tissue for markers of neocortical GABAergic interneuron subtypes—Parvalbumin (PV) and Calretinin (**Figure 3C/D**). We found presence of both FLAG^+^/PV^+^ and FLAG^+^/Calretinin^+^ cells, indicating both serotypes readily transduce these GABAergic interneuron populations.

Another manifestation of the irregular transduction pattern was the presence of FLAG^+^ neurons in large clusters. The presence of these clusters was correlated with areas of high transduction, such as the somatosensory or visual cortex (**Figure 2 A2/B2, Figure 3E**). We also noted a higher proportion of clusters in the cortex and often near large descending blood vessels (**Figure 3E’**). To examine this further, we co-stained the tissue for GLUT1, a marker in endothelial cells that line the blood-brain barrier (**Figure 3F/F’**).

While many clusters were found surrounding or neighboring medium to large vessels, this association was not exclusive. We also examined whether a high concentration of transgene expressing cells can result in local cellular pathology. We stained the tissue for IBA1, a microglia marker, but no signs of microglial activation were observed near these clusters (**Figure 3G1/2**). Furthermore, we did not observe any signs of neuronal loss, gliosis, or astrocyte activation, as assessed by GFAP co-staining (data not shown).

We further assessed neuronal transduction and transgene expression resulting from AAV-PHP.eB and AAV9 ICV administration by quantifying FLAG^+^ and NeuN^+^ cells in matched cortical regions along the anterior-posterior axis of the brain, namely in the prefrontal cortex (vmPFC), primary motor cortex (M1), somatosensory cortices (S1/S2), posterior parietal cortex (PPC), and visual cortex (V1/V2) (see methods for details). Overall, 336 ROIs (∼1.4 million NeuN^+^ cells) and 343 ROIs (∼1.3 million NeuN^+^ cells) were quantified in both the AAV-PHP.eB and AAV9 groups, respectively. Consistent with qualitative observations, the AAV-PHP.eB cohort displayed more expression in all analyzed brain regions compared to the AAV9 group (**Figure 4A**). There was no statistical difference between regions along the anterior-posterior axis in the PHP.eB group (combined ipsilateral and contralateral sides), with median % neuronal labeling as follows: PFC=8.96% (n=59), M1=10.39% (n=69), S1/S2=11.99% (n=72), PPC=10.95% (n=70), V1/V2=8.03% (n=66), P = 0.286. In contrast, in the AAV9 cohort, there was a significant difference between areas (P = 0.003), with visual cortex showing a higher % of labeled neurons than M1, S1/S2 and PPC. Median values were as follows: PFC=2.14% (n=66), M1=1.88% (n=66), S1/S2=1.57% (n=69), PPC=1.93% (n=65), V1/V2=4.22% (n=77). When comparing the transduction rate between the hemispheres, no significant differences were found in either group: PHP.eB ipsilateral 10.83% (n=169) vs. contralateral 11.87% (n=167) (P = 0.348), and AAV9 ipsilateral 1.90% (n=171) vs. contralateral 2.46% (n=172) (P = 0.299) (**Figure 4B**). Importantly, there was a significant difference in cortical neurons transduction rate between the two serotypes, with median PHP.eB value of 11.1% (mean = 12.1 ± 0.72) compared to 2.2% (mean = 3.1 ± 0.30) for AAV9 (P < 0.001) (**Figure 4C**).

**Figure 4.**
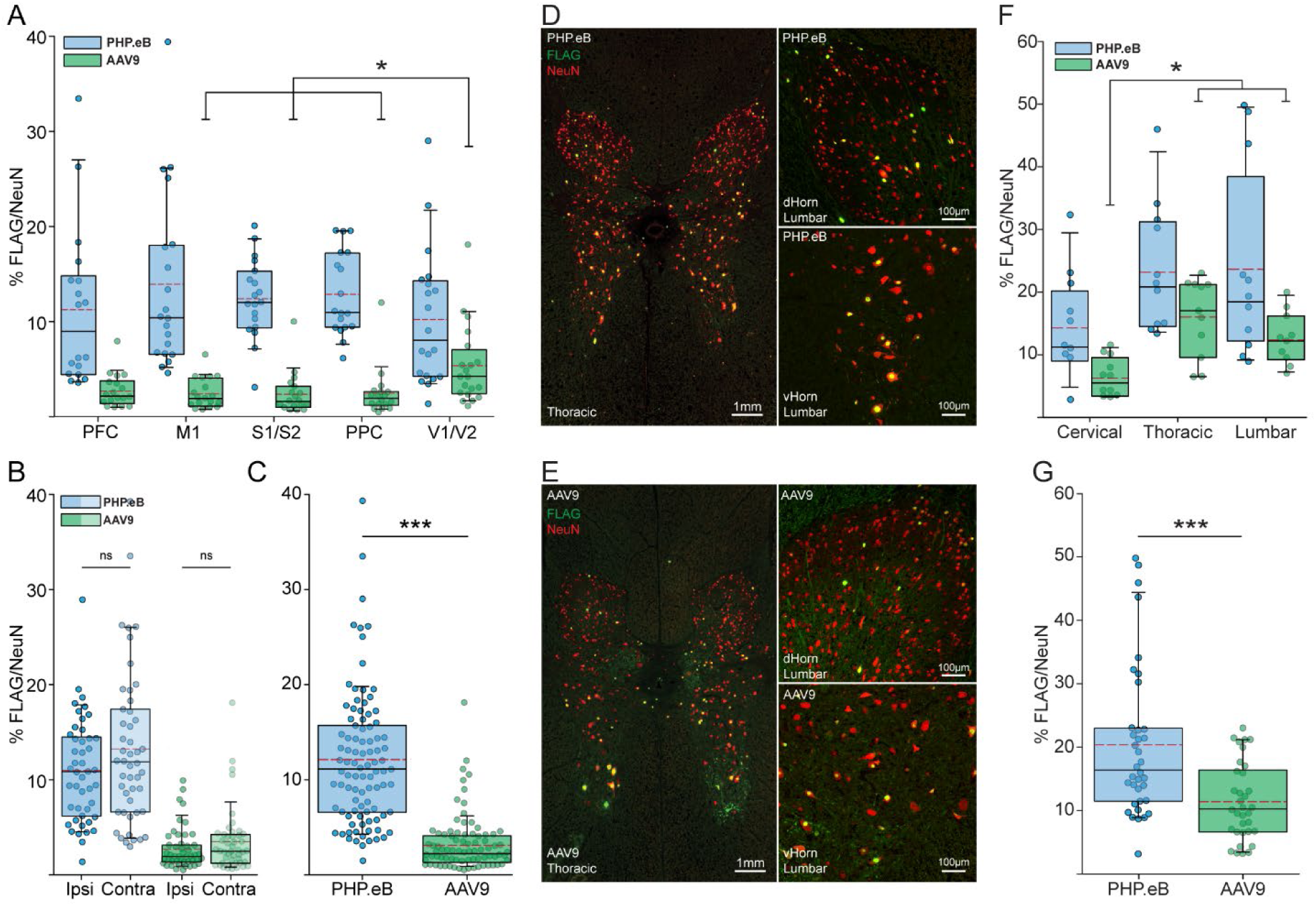
AAV-PHP.eB yields significantly higher neuronal transduction rates in the NHP cortex and spinal cord compared to AAV9. Quantification of transduced neurons in defined cortical and spinal areas; the rate of transduction was measured as the fraction of FLAG^+^ to NeuN^+^ cells (%) and is shown in box-and-whisker plots (dashed red line represents mean value, individual measures are shown as dots and overlayed). **A)** Comparison of transduction rates in matched anatomical locations along the anterior-posterior axis of the brain. One-way ANOVA on Ranks was used for within-cohort comparisons (PHP.eB: P = 0.286; AAV9: P = 0.003), followed by All Pairwise Multiple Comparison Procedures (Dunn’s Method) for AAV9 (P < 0.05). **B)** Comparison of transduction rates between ipsilateral and contralateral sides of the injection. Mann-Whitney Rank Sum Test was used (PHP.eB: P = 0.348; AAV9: P = 0.299). **C)** Brain-wide transduction rates for PHP.eB and AAV9 groups (P < 0.001, Mann-Whitney Rank Sum Test). **D/E:** Typical expression patterns in the spinal cord for PHP.eB **(D)** and AAV9 **(E)** at the thoracic level; zoomed in micrographs on the right illustrate neuronal transduction of the dorsal and ventral horn in the lumbar segments of the cord. Note robust labeling of large lower motor neurons. A pseudo-coloring scheme and scale bars are shown. **F)** Rate of transduction for PHP.eB and AAV9 cohorts in cervical, thoracic, and lumbar segments of the spine. Comparisons within cohorts were performed using One-way ANOVA on Ranks (P = 0.063 for PHP.eB group, P < 0.001 for AAV9) followed by All Pairwise Multiple Comparison Procedures (Dunn’s Method). **G)** Comparison of the rate of transduction for the whole spinal cord in PHP.eB and AAV9 groups (P < 0.001, Mann-Whitney Rank Sum Test). **Abbreviations:** PFC – prefrontal cortex, M1 – primary motor cortex, S1/S2 –somatosensory cortex area 1 and 2, PPC – posterior parietal cortex, V1/V2 – visual cortex areas 1 and 2, Ipsi – ipsilateral hemisphere, Contra – contralateral hemisphere.

Next, we compared AAV9 and AAV-PHP.eB expression in the spinal cord. Cervical, thoracic, and lumbar segments of the spine were co-stained for FLAG and NeuN and quantified (**Figure 4D-G**). Both groups transduced all levels of the spinal cord, with FLAG^+^ cells detected in all regions of the gray matter (dorsal and ventral horns, as well as the peri-central canal area) (**Figure 4D/E**). Higher percentage of neuronal labeling was observed in the more distal parts of the cord; median PHP.eB - cervical = 11.22% (n = 12), thoracic = 20.83% (n = 12), lumbar = 18.46% (n = 12); median AAV9-cervical = 5.48% (n = 12), thoracic = 17.01% (n = 11), lumbar = 12.18% (n = 11). (**Figure 4F**). When considering all segments together, AAV-PHP.eB had ∼1.5-fold higher neuronal transduction in the spinal cord compared to AAV9: median rate for AAV-PHP.eB 16.39% (n = 36) vs. 10.26% (n = 34) for AAV9 (**Figure 4G**).

To complement the histological examination, we also sampled selected brain areas and peripheral organs for viral DNA by ddPCR (**Figure 5**). In the motor cortex, AAV-PHP.eB group exhibited a median value of 5.1 vector genome copies/diploid genome (copies/dg), whereas AAV9 group totaled 0.77 copies/dg (P < 0.001) (5.5-fold difference with IHC vs. 6.6 with ddPCR). Similarly, in the visual cortex, the AAV-PHP.eB group had a median of 2.55 copies/dg compared to 1.18 copies/dg for AAV9 (P < 0.001) (1.9-fold increase assessed by IHC vs. 2.16 with ddPCR). In agreement with histological observations, both vectors exhibited minimal penetration of the putamen, with no significant difference between the groups (0.12 copies/dg for AAV-PHP.eB vs. 0.13 copies/dg for AAV9; P = 0.053). The analysis of the peripheral tissues revealed that much of the vector leaks into the general circulation and gets distributed though the body. Vector concentration was relatively low in the skeletal muscle (PHP.eB: 0.40 copies/dg vs. 0.29 copies/dg for AAV9; P < 0.001) and the kidney (PHP.eB: 0.43 copies/dg vs. 0.46 copies/dg for AAV9; P = 0.66), moderate in the heart (PHP.eB: 1.02 copies/dg vs. 1.77 copies/dg for AAV9; P = 0.034), and notably high in the liver (PHP.eB: 188.6 copies/dg vs. 234.3 copies/dg for AAV9; P = 0.06).

**Figure 5.**
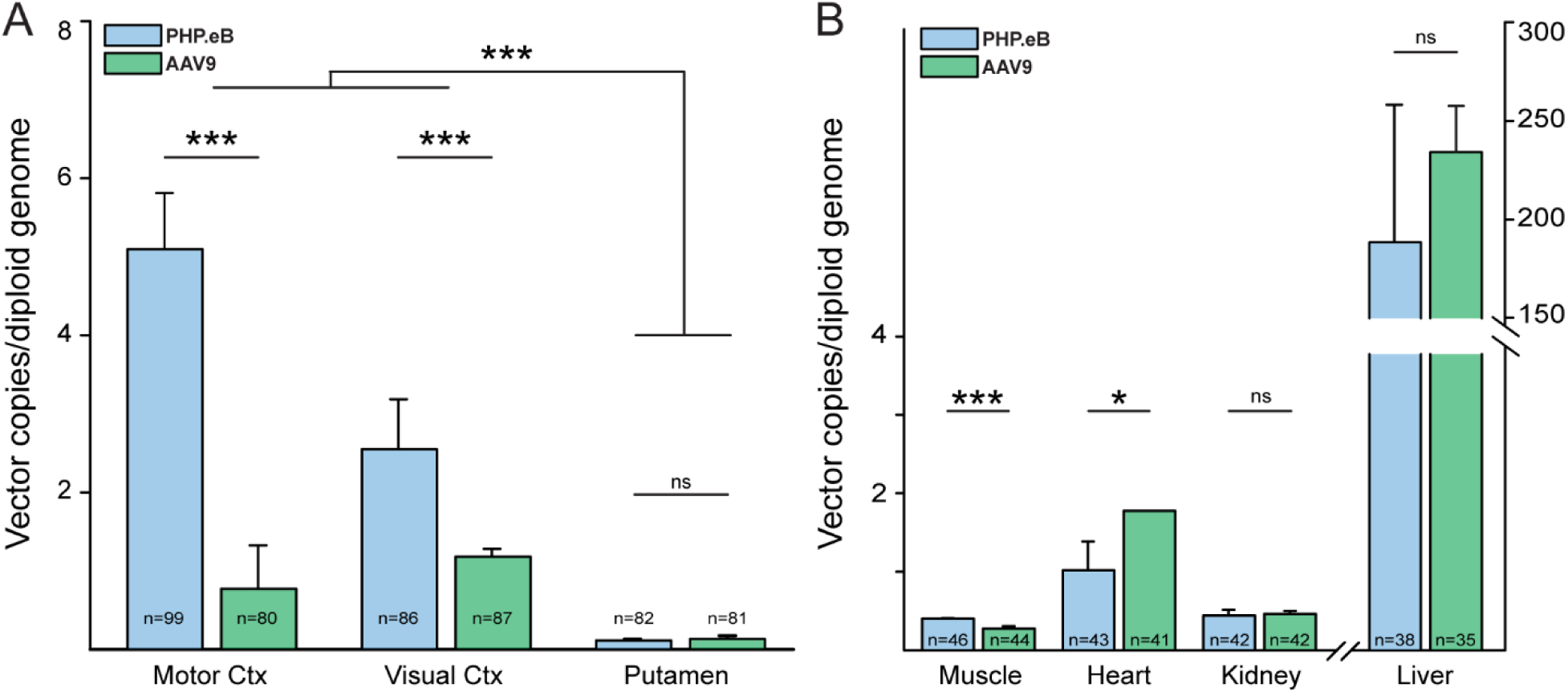
Viral DNA detection in the brain and peripheral tissues – a ddPCR analysis. AAV-PHP.eB yields a significantly higher rate of transduction in the NHP cortex compared to AAV9. The vector copy number per cell was calculated relative to the MRFAP1 gene and is presented in bar graphs (median ± SE). **A:** Comparison of matched brain areas for PHP.eB and AAV9 groups. PHP.eB demonstrated higher transduction rates than AAV9 in the motor (median 5.1 vs. 0.77 copies/cell) and visual cortex (median 2.55 vs. 1.18 copies/cell) (P < 0.001, Mann-Whitney Rank Sum Test). In the putamen, both vectors showed very little penetration, with no significant difference between groups (median 0.12 vs. 0.13 copies/cell; P = 0.053, Mann-Whitney U). **B:** Majority of the vector leaks into peripheral tissues. Vector concentration in selected peripheral tissues: skeletal muscle (gastrocnemius) PHP.eB: 0.40 vs. AAV9: 0.29 (P < 0.001), heart (ventricle) PHP.eB: 1.02 vs. AAV9: 1.77 (P = 0.034), kidney (cortex) PHP.eB: 0.43 vs. AAV9: 0.46 (P = 0.66), and liver PHP.eB: 188.6 vs. AAV9: 234.3 (P = 0.06) (note different scale on the right y-axis for the liver).

## V. Discussion

This study provides a comprehensive comparison of neuronal transduction efficiency and vector biodistribution of AAV-PHP.eB versus AAV9 following unilateral ICV administration in juvenile macaques. Using a neuron-specific promoter and a nuclear-localized reporter gene, we systematically assessed transduction across cortical, subcortical, spinal, and peripheral tissues. While both vectors achieved widespread CNS distribution with similar expression patterns, AAV-PHP.eB exhibited significantly higher percentage of neuronal labeling throughout the brain and spinal cord. These findings highlight the superior efficiency of AAV-PHP.eB and its potential advantages in CNS-targeted gene therapy.

### Key Findings

Histological and molecular analyses revealed that AAV-PHP.eB outperformed AAV9 in most assessed brain regions. In cortical areas, AAV-PHP.eB achieved transduction rates six times higher in motor and somatosensory cortices, with an average four-fold increase across the cortex. In the spinal cord, the magnitude of this difference was not as pronounced, with AAV-PHP.eB averaging ∼1.5-fold increase in neuronal labeling rate compared to AAV9. In contrast, both vectors exhibited minimal transduction in deeper brain structures, highlighting persistent challenges in achieving efficient subcortical delivery with CSF-based routes in NHPs (19–22).

Although AAV-PHP.eB’s enhanced BBB-crossing and improved performance in rodent models is attributed to binding to LY6A receptor, which is absent in NHPs, AAV-PHP.eB still showed superior transduction when delivered via ICV route (8,12,22). Interestingly, in mice, ICV administration of AAV-PHP.eB showed no transduction or tropism advantage over AAV9 in either neonates or adults (12). Chatterjee et al. reported enhanced cortical and cerebellar transduction with AAV-PHP.eB over AAV9 following ICM delivery in adult rats (23). The only other study evaluating AAV-PHP.eB’s transduction efficiency in NHPs was Arotcarena et al. (2021), who compared it with its precursor, AAV-PHP.B, in adult rhesus macaques following a single lumbar intrathecal injection (13). AAV-PHP.eB demonstrated significantly greater and broader CNS transduction than AAV-PHP.B. This suggests that mechanisms beyond BBB-crossing influence AAV-PHP.eB’s enhanced brain penetration and transduction efficacy compared to its predecessors, warranting further investigation.

An important observation in this study was the uneven, patchy distribution of transgene expressing neurons within the cortex, an expression pattern reported before in NHPs (6,10,21). Areas of dense neuronal labeling were often adjacent to regions with sparse transduction, indicating non-uniform viral particle distribution post-injection. In highly transduced regions prominent clustering of labeled cells was frequently observed. The association of these clusters with brain vasculature suggests that penetrance of viral particles deep into the parenchyma is mediated by CSF flow along perivascular spaces, commonly referred to as the glymphatic pathway (24). A critical observation in this respect is that clusters were observed along some, but not all, vessels, indicating that the direction of CSF flow along the glymphatic spaces and viral clearance mechanism might be key factors influencing the spread and extent of transduction following direct CSF administration of AAVs.

The extensive histological evaluation and systematic quantification of neuronal transduction across multiple brain areas in this study is noteworthy. Many studies rely on ubiquitous promoters, such as CMV or CAG, and cytosolic markers like EGFP, which visualize transduced cell populations but hinder accurate quantification due to intense neuropil labeling (6,13,22). Consequently, NHP studies often resort to qualitative or indirect methods for reporter gene quantification (e.g., bulk tissue qPCR/ddPCR or fluorescent signal intensity quantification). Here, the neuron-specific hSyn1 promoter and nuclear-localized reporter enabled precise quantitative assessment of neuronal transduction.

### Implications for gene therapy

The higher transduction efficiency of AAV-PHP.eB suggests potential utility as a more effective vector for gene therapies targeting widespread cortical neuronal transduction. However, limited transduction in subcortical regions, which are critical for addressing movement disorders like Parkinson’s or Huntington’s disease, remains a challenge. Future studies should explore alternative delivery methods or capsid modifications to improve transduction in these deep brain regions. Ongoing capsid engineering may yield variants with enhanced BBB-crossing capabilities in NHPs, enabling more uniform distribution via brain circulation (2,8,25–27).

### Potential Limitations and Challenges

Despite equal dosing per kilogram body weight (4 × 10¹³ vg/kg), differences in cohort body weight may have introduced bias, potentially underestimating AAV9 transduction efficiency. Additionally, while the neuron-specific promoter and nuclear-localized reporter enabled precise quantification, the exclusion of non-neuronal cell populations, (e.g., astrocytes, oligodendrocytes), limits the broader applicability of these findings.

### Conclusion Future Directions

AAV-PHP.eB achieves significantly higher neuronal transduction efficiency than AAV9 in NHPs following ICV administration, making it a promising candidate for CNS gene therapy, particularly for cases requiring extensive cortical neuron transduction. Limited subcortical transduction and peripheral leakage remain significant challenges. Future research should focus on optimizing delivery routes and vector designs to enhance deep brain targeting and minimize off-target effects. These findings provide a strong foundation for refining AAV-based gene therapies for CNS disorders.

## VI. Data Availability

Data from this study will be made available upon request.

## Acknowledgments

We thank members of the Washington National Primate Research Center for providing excellent husbandry animal care, surgical support, and assistance with tissue collection for the study, in particular Christopher English, Britni Curtis, Jesse Day, and Sandi Thelen. We thank Bargavi Thyagarajan for support managing WaNPRC master services agreements. We acknowledge support to WaNPRC by the NIH Office of Research Infrastructure Programs (ORIP) under award number P51OD010425 and U420D011123. We thank the University of Washington Diagnostic Imaging Sciences Center (DISC) Magnetic Resonance Research Laboratory for support with macaque MRI.

## VII. Author Contributions

M.G.F., A.H., K.G., B.L., E.L., and J.T.T. conceived the experiments, M.G.F, L.N., N.T., N.W., A.H., and M.B. conducted experiments, and M.G.F., L.N., N.T., and J.T.T. analyzed results. All authors reviewed the manuscript.

## VIII. Funding

The work was funded by the Allen Institute for Brain Science.

## IX. Ethical Approval

All work with macaque monkeys conformed to the guidelines provided by the US National Institutes of and was approved by the University of Washington Institutional Animal Care and Use Committee (IACUC).

## X. Competing Interests

The authors declare no competing interests.

## Notes

### Competing Interest Statement

The authors have declared no competing interest.

